# Metacommunity analyses show increase in ecological specialisation throughout the Ediacaran

**DOI:** 10.1101/2021.05.17.444444

**Authors:** Rebecca Eden, Andrea Manica, Emily G. Mitchell

## Abstract

The first animals appear during the late Ediacaran (572 – 541 Ma); an initial diversity increase was followed by a drop, interpreted as catastrophic mass extinction. We investigate the processes underlying these changes using the “Elements of Metacommunity Structure” framework. The oldest metacommunity was characterized by taxa with wide environmental tolerances, and limited specialisation and inter-taxa interactions. Structuring increased in the middle metacommunity, with groups of taxa sharing synchronous responses to environmental gradients, aggregating into distinct communities. This pattern strengthened in the youngest metacommunity, with communities showing strong environmental segregation and depth structure. Thus, metacommunity structure increased in complexity, with increased specialisation and resulting competitive exclusion, not a catastrophic environmental disaster, leading to diversity loss in the terminal Ediacaran, revealing that the complex eco-evolutionary dynamics associated with Cambrian diversification were established in the Ediacaran.

## Introduction

One of the most dramatic events in the history of Earth is the sudden appearance of animals in the fossil record during the Ediacaran period (635-541 Ma), after billions of years of microbial life (5,6). Ediacaran anatomies are particularly difficult to compare to modern phyla, which has hampered our understanding of Ediacaran evolution and how Ediacaran organisms relate to the Cambrian Explosion and extant animal phyla (7). Patterns of taxonomic, morphological and ecospace diversity change dramatically during the Ediacaran (8,9), which has led to the suggestion of several evolutionary radiations, corresponding to the Avalon, White Sea and Nama assemblages (5,10–12). These three assemblages are formed by the taxonomic groupings of communities which occupy partially overlapping temporal intervals and water-depths, with no significant litho-taphonomic or biogeographic influence (10,11,13). The oldest assemblage, the Avalon (575-565 Ma), exhibits relatively limited ecological and morphological diversity (8,9), with only limited palaeoenvironmental influence on its composition and taxa interactions (14–17). The White Sea assemblage (558-550 Ma) shows a large increase in morphological diversity, including putative bilaterians, in tandem with a greater ecological diversity (8) and environmental influence (18). The Nama assemblage (549-543 Ma) includes the oldest biomineralizing taxa, and records a decrease in taxonomic diversity (8,19–21). The reduction in taxonomic diversity and community composition complexity between the White Sea and Nama has been suggested to correspond to a post-White Sea extinction around 550 Ma, which eliminated the majority of Ediacaran soft-bodied organisms (12,22–24).

Previous studies have focused primarily on defining the different assemblages and the underlying causes for the different assemblages (10,11,13), looking at taxonomic and morphological diversity between assemblages (8,25) with little investigation of how the ecological structure within the assemblages differs. The network structures of Ediacaran body-fossil and trace fossil taxa were compared by Muscente et al. (12) using modularity to compare found networks (and sub-networks which correspond to assemblages) to random networks. However, these prior cluster and network analyses have not assessed the relative frequency of taxa co-occurrences within assemblages; i.e. whether they are were statistically different to what would be expected by random chance, nor compared the ecological structure within each assemblage to known ecological models.

In this study, we will investigate the structuring mechanisms within these assemblages using three analyses which have not been previous used to investigate Ediacaran macro-ecology. We used a binary presence-absence matrix for 86 Ediacaran localities and 124 taxa, with paleoenvironment, depth, lithology, time and assemblage data from (11,24) (Fig. S1). First, we will use the “Elements of Metacommunity Structure” (EMS) framework to investigate emergent properties of groups of connected communities that may arise from taxa interactions, dispersal, environmental filtering and the interaction of these factors (2–4) (Fig. 1). The EMS framework is a hierarchical analysis which identifies properties in site-by-taxa presence/absence matrices which are related to the underlying processes shaping taxa distributions (4), but to date has limited application to the fossil record (26). Three metacommunity metrics are calculated to determine the structure: Coherence, Turnover, and Boundary Clumping (2–4). The values of these metrics determine where the metacommunity fits within the EMS framework (Fig. 1), with different metric combinations indicating different underlying processes behind the metacommunity structure. Secondly, we will test to determine which pair-wise taxa co-occurrences are significantly non-random, and whether any non-random co-occurrences are positive or negative. Finally, we use reciprocal averaging to ordinate the sites based on their community composition and then correlate the first-axis ranking with depth to test whether within-assemblage community composition is correlated with depth.

**Figure 1:**
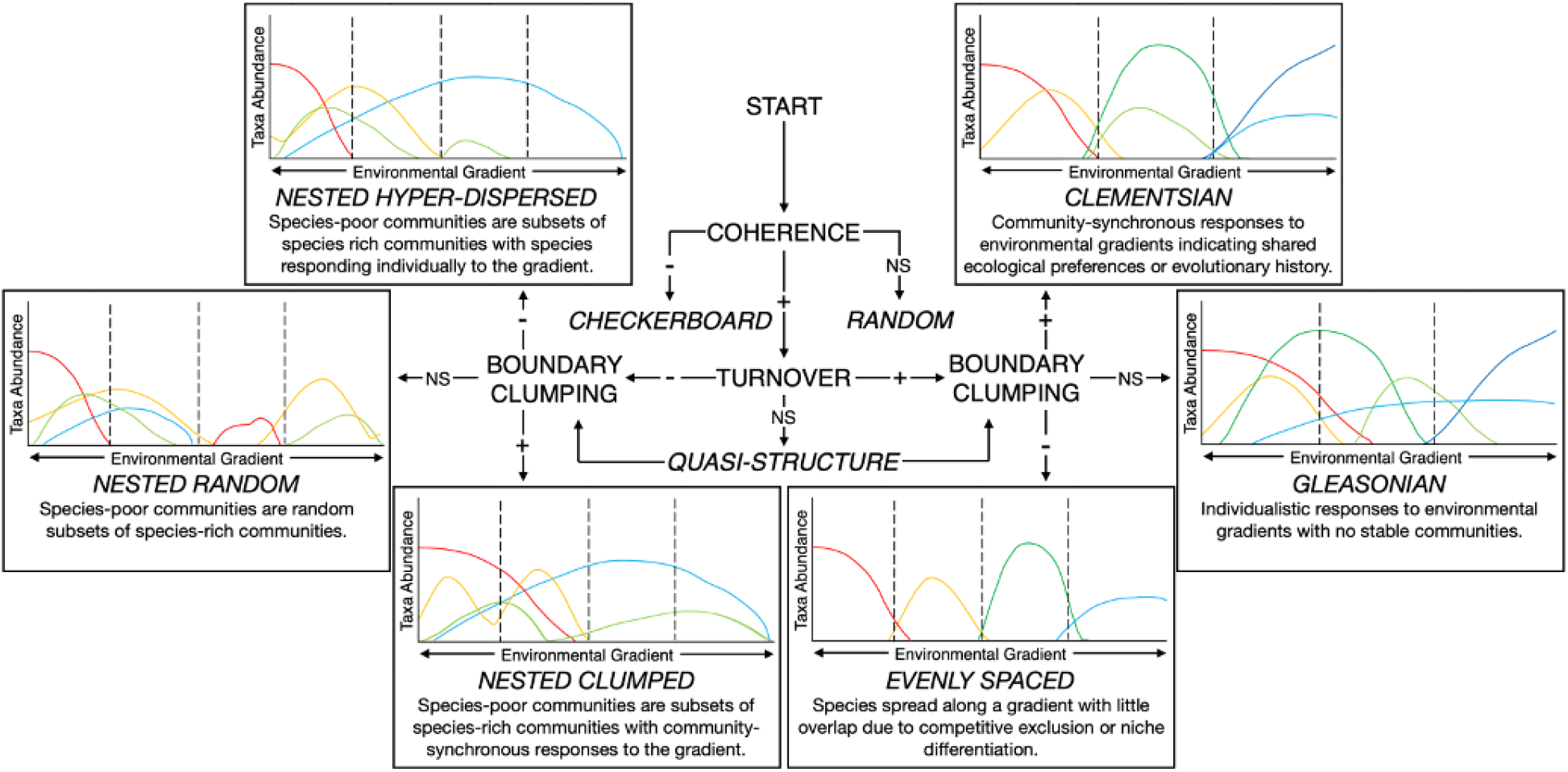
Idealized metacommunity structures. These graphs show taxa abundance patterns in idealized metacommunities of several taxa (represented by different colors) which respond to a latent environmental gradient (they exhibit significant positive Coherence).

## Results

First, we pooled all the sites irrespective of their assemblage, in order to test whether the assemblage definitions harbored distinct communities (Fig. 2). When analysing the total dataset, we found positive Coherence, Turnover and Boundary Clumping, characteristic of a Clementsian structured metacommunity (Fig. 3a, Table 1). Site ordination scores were significantly correlated with assemblage (*H*=57.686, *df*=2, *p*<0.001, Appendix Table 4) and depth (*H*=51.649, *df*=7, *p*<0.001, Table S3), indicating that the structuring in the dataset was due to differences in depth or other assemblage-specific characteristics. To further investigate the nature of this structuring, we focused on the pair-wise co-occurrence patterns: virtually all positive associations resulted from taxa specific to the same assemblage (96.8% of positive associations), and all negative associations from taxa found in different assemblages (100% of negative associations). Therefore, our analyses are consistent with previous studies (11,12) in finding that Ediacaran taxa are highly segregated by assemblage, as well as confirming a role for depth (Table 2) in structuring the assemblages (10,11). At this broad level of analysis, the strong assemblage signal, at least partially dependent on depth specialisations, obscures any other biotic or abiotic pattern.

**Table 1:** Metacommunity analyses. Metacommunity values for Coherence, Turnover and Boundary Clumping with interpretation of metacommunity structure within the Elements of Metacommunity Structure framework. Z is the Z-score, p is the p-value, simMean is the simulated mean value of the metric. Significant p-values are highlighted in bold.

|  | Coherence |  |  |  | Turnover |  |  |  | Boundary Clumping |  |  | Interpretation |
| --- | --- | --- | --- | --- | --- | --- | --- | --- | --- | --- | --- | --- |
| Group | Coherence | p | simMean |  | Turnover | p | simMean |  | Moritsita's Index | p |  |  |
| Total | 2,230 | **<0.001** | 6,690 | + | 1,270,000 | **<0.001** | 1,010,000 | + | 6.390 | **<<0.001** | + | Clementsian |
| Avalon | 173 | **<0.001** | 309 | + | 5,120 | **0.007** | 7,650 | - | 1.880 | **<0.001** | + | Clumped species loss (nested subsets) |
| Avalon (Margin slope) | 167 | **<0.001** | 221 | + | 1,890 | **0.007** | 3,580 | - | 1.980 | **<0.001** | + | Clumped species loss (nested subsets) |
| Avalon (Outer shelf) | 10 | 0.510 | 11 | + | 0 | 0.119 | 21 | - | 1.330 | 0.156 | + | No significant metacommunity structure |
| White Sea | 695 | **<0.001** | 1,340 | + | 61,900 | 0.098 | 48,200 | + | 3.020 | **<0.001** | + | Clementsian Quasi Structure |
| White Sea (Deep subtidal) | 60 | **<0.001** | 217 | + | 2,780 | 0.230 | 2,030 | + | 3.450 | **<0.001** | + | Clementsian Quasi Structure |
| White Sea (Middle shelf) | 182 | **<0.001** | 344 | + | 5,300 | 0.110 | 3,580 | + | 4.700 | **<0.001** | + | Clementsian Quasi Structure |
| White Sea (Outer shelf) | 0 | **0.004** | 16 | + | 66 | **0.044** | 29 | + | <0.001 | 0.227 | - | Gleasonian |
| Nama | 21 | **<0.001** | 133 | + | 688 | 0.435 | 640 | + | 2.350 | **<0.001** | + | Clementsian Quasi Structure |

**Table 2:** Reciprocal averaging analyses.

|  | Rs | Kruskal-Wallis | df | p-value |
| --- | --- | --- | --- | --- |
| All assemblages site scores x assemblage | - | 57.686 | 2 | **<0.001** |
| All assemblages site scores x depth | - | 51.649 | 7 | **<0.001** |
| Depth x assemblage | - | 53.987 | 2 | **<0.001** |
| Avalon site scores x depth | -0.360 | - | - | 0.051 |
| White Sea site scores x depth | 0.014 | - | - | 0.945 |
| Nama site scores x depth | -0.728 | - | - | **0.007** |

Spearman's rank correlations (for continuous variables) and Kruskal-Wallis tests (for discrete variables) between site scores ( $R_s$ ) of each dataset obtained from reciprocal averaging and a variable, either assemblage or depth. Significant p-values are highlighted in bold.

**Figure 2:**
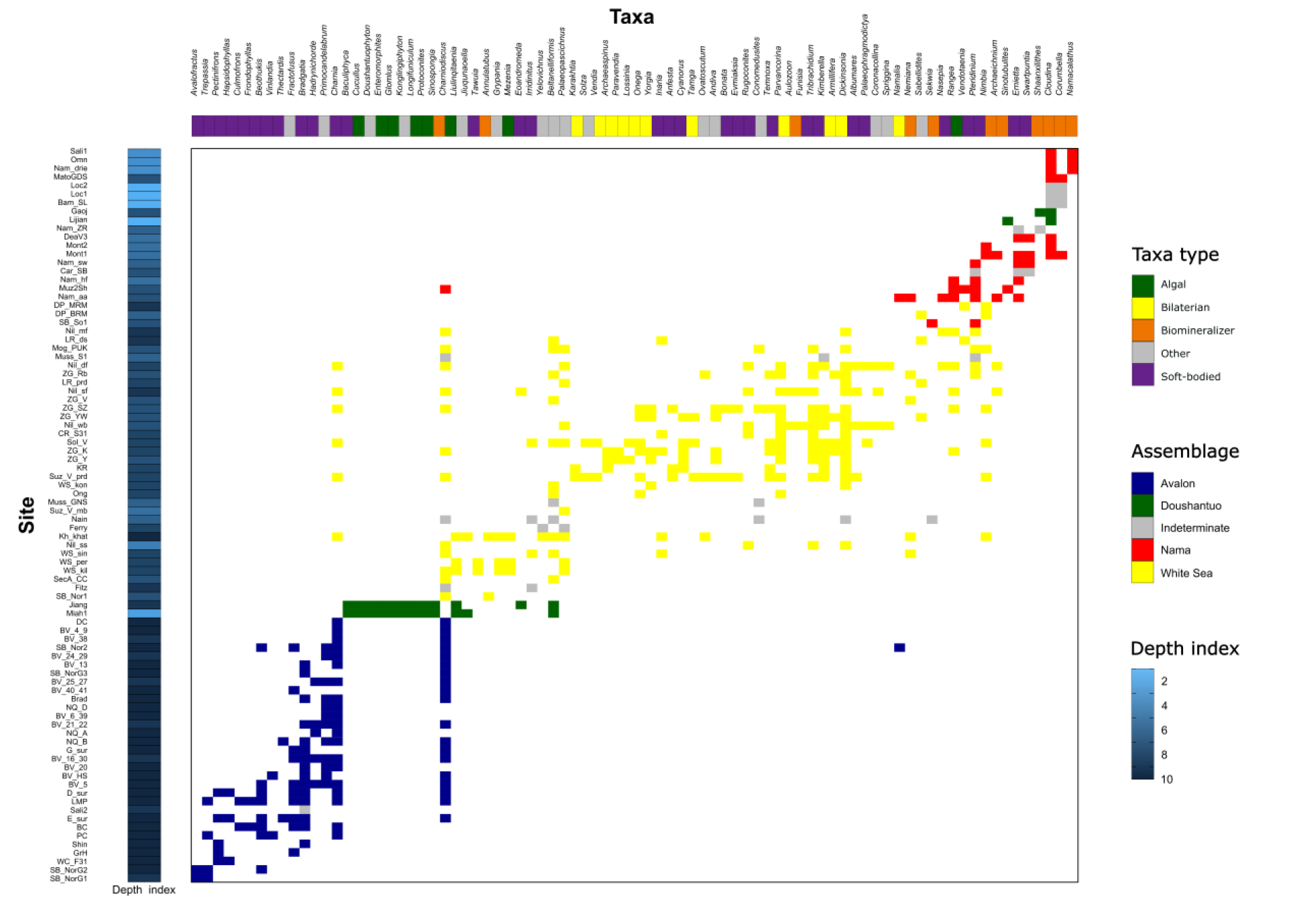
Ordinated data table. The assemblage and palaeoenvironment for each locality are given on the left. Right plot shows the incidence matrix for taxa (columns) for all sites (rows) along the inferred environmental gradient after ordination. Ordination was calculated according to occurrence resulting from the overall metacommunity analysis. The presence of a taxon is given by a colored square, absence by white. Doushantuo and Indeterminate sites were excluded from the assemblage-level analyses. Depth index indicates the relative depth of the locality cf. Boag et al. 2016, as determined by palaeoenvironment.

**Figure 3.**
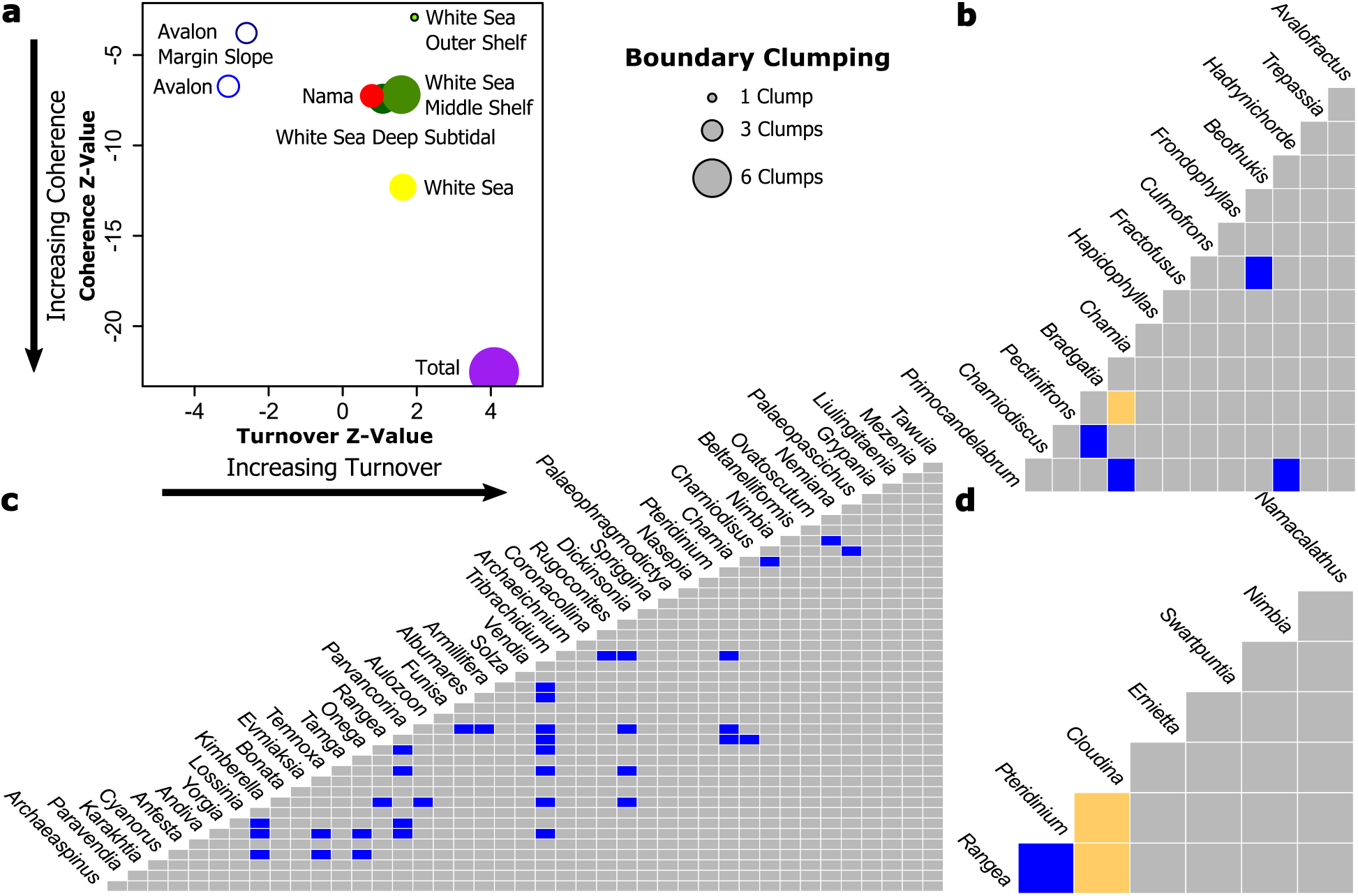
Metacommunity analyses and co-occurrence matrices for each assemblage. **a**, Metacommunity plot shows a summary of the metacommunity analyses. Nested species loss is shown by an open circle; Clementsian by a closed circle; Gleasonian by a black ringed circle. The Avalonian outer shelf had a random structure and so is not shown. Co-occurrence matrices for **b**, the Avalon metacommunity, **c**, the White Sea metacommunity and **d**, the Nama metacommunities. Positive associations are blue, negative associations are yellow.

We then narrowed our level of analysis by focusing on each assemblage in turn. The Avalonian metacommunity displays significant positive Coherence and Boundary Clumping, but significant negative Turnover (Fig 2, Table 1), characteristic of a pattern of “clumped species loss” (1,3,27). Site ordination scores were not significantly correlated with depth (*Rs=*-0.360, *p=*0.051, Table S2). The metacommunity structure was the same for the Avalonian margin slope metacommunity, but the outer shelf showed no significant structuring (Figs. 2, Fig. S3, Table S, Tables S2, S3). The predominance of positive over negative taxa co-occurrences is consistent with previous studies which found little evidence for lateral resource competition between Avalonian taxa (14,18). The lack of depth and palaeoenvironmental correlation with metacommunity structure supports suggestions that Avalonian organisms have the widest niches and lowest provinciality amongst the Ediacaran biotas (10,15).

The Avalonian metacommunity displays a structure of “clumped species loss”, whereby taxa-poor communities form nested subsets of increasingly taxa-rich communities, with predictable patterns of taxa loss associated with variation in taxa characteristics (3). Differences in Avalonian community composition have been suggested to represent different stages of community succession (16), a process which would result in the observed pattern of clumped taxa loss with early and late succession communities forming less-diverse nested subsets of maximally diverse mid-succession communities. However, Avalonian metacommunity structure had not previously been statistically compared to multiple different models (here 12), and so analytically compared to different hypotheses. Our results find that the Avalonian metacommunity exhibits clumped taxa loss which supports this successional model hypothesis. This hypothesis is further supported by the co-occurrence analyses which found a negative association between late-succession and early-stage taxa (*Charnia* and *Pectinifrons*), and positive associations between late (*Primocandelabrum* and *Charnia*), middle (*Bradgatia* and *Charniodiscus*) and early (*Beothukis* and *Fractofusus*) stage taxa (Fig. S2. Table S3, Supplementary Discussion) (16). The taxa associations of the Avalonian sites in the UK form a subset of the associations from the Canadian sites, suggesting structuring is not due to geographic or abiotic differences between the two areas (Fig. S3. Table S3, Supplementary Discussion). Therefore, observed differences in community composition between UK and Canadian Avalonian sites may reflect which stages in community development are preserved, and thus represent a biotic rather than a geographic signal.

The White Sea metacommunity displays significant positive Coherence and Boundary Clumping and non-significant positive Turnover, characteristic of a “Clementsian quasi-structure” (3) with no correlation between site ordination scores and depth (*Rs=*0.014, *p=*0.945, Fig. 2; Tables 1-2).

Metacommunity structure within the White Sea assemblage differed with palaeoenvironment: the deep subtidal and middle shelf metacommunities displayed “Clementsian quasi-structures”, while the outer shelf metacommunity was characterized by a Gleasonian structure, with significant positive Coherence and Turnover and non-significant Boundary Clumping (Fig. 3, Fig. S4. Table 1). Co-occurrence analyses found that 11 of the 32 positive associations found for the whole assemblage were preserved when focusing on the palaeoenvironmental subsets (Fig. 3, Fig. S4. Tables S 6-7, Supplementary Discussion). Because each positive association is only determined to be a positive association only when it occurs significantly more than expected by random, if the taxa were behaving exactly the same way in each assemblage sub-set, we would expect the same positive associations to occur in each palaeoenvironmental subset, i.e. the null hypothesis would be 32/32 positive associations. Our finding that 11/32 illustrated positive associations is significantly different from this null hypothesis; therefore the community interactions are significantly different between the subsets and the assemblage as a whole. Several taxa seemed to behave differently in the different palaeoenvironments, with unique taxa associations in each subdivision despite very similar community composition. (Fig. S4. Tables S6-7, Supplementary Discussion). Variation in both inter-taxa interactions and environmental factors can cause organisms to exhibit behavioral plasticity, acting differently in different environments. Differences in behavior in the same taxon may reflect the inclusion of several taxa with different environmental preferences and behaviors within one taxonomic group (e.g. *Dickinsonia*). However, as most Ediacaran taxonomic groupings are monotypic the differences in the behavior of the taxa in different environments is more likely due to plastic responses to, for example, variations in resource limitation or the presence of different competitors or ecosystem engineers.

The majority of taxa pairwise associations found in the Russian White Sea are also found in the pooled White Sea metacommunity (24/37) However, 13 of the pooled associations are only present when only analysing Russian localities, suggesting that some of the structure in the dataset may be due to geography (Tables S5, S8). There is marked geographic variation in community composition between the Russian and Australian White Sea localities (24), so the non-shared associations may reflect a greater endemism within the White Sea assemblage compared to the Avalonian assemblage, where the UK sites formed a perfect subset of the Canadian sites

The Nama metacommunity has significant positive Coherence and Boundary Clumping and non-significant positive Turnover and so displays a “Clementsian quasi-structure” (Table 1) (3). Unlike the Avalon and White Sea sites, ordination scores were significantly correlated with depth (*Rs=-*0.728, *p=*0.007, Table 2) and so the Clementsian structuring occurs along a depth gradient. Pairwise taxa co-occurrences revealed significant negative associations between biomineralizers and soft-bodied taxa, and a significant positive association between two soft-bodied taxa (Fig. 3, Table S9). There were more negative than positive associations (Fig. 3, Table S9). These results statistically confirm previous observations of separation between biomineralizers (e.g. *Cloudina*) and soft-bodied organisms (such as *Pteridinium* and *Rangea*).

These patterns of segregation are unlikely to be purely facies-based control for our data set. In the Nama assemblage there are both deep and shallow water carbonate facies. The deep water Nama facies (Dengying Fm) has a more similar community composition to the other deep water Nama sites than to the carbonate sites and thus help to strengthen the pattern of taxa segregation by depth as opposed to counteract it (which is what we would expect if there was purely facies-based control of taxa separation). If the habitat specialization was a reflection on biomineralization alone, that we would expect to see the biomineralizes behave in broadly similar patterns, and the soft-bodied taxa to also behave similarly to each other. Of the seven taxa that were sufficiently abundant to be included in these analyses, one was a putative microbial colony (*Nimbia* (28)), two were biomineralisers (*Cloudina* and *Namacalathus*) and the remaining four were soft-bodied taxa (*Rangea, Pteridinium, Ernietta* and *Swartpuntia*). *Nimbia* did not show any significant associations with the other taxa, although (like *Namacalathus*) it was only present in 2 sites, this lack of associations may be due in part to to small sample sizes. The two biomineralisers behaved in different ways – while *Cloudina* showed negative associations with the soft-bodied Rangea and Pteridium. *Namacalathus* did not show any significant associations with any taxa, but could be biased by the number of sites in which it is present. The soft-bodied taxa also did not behave in a uniform way, with *Ernietta* and *Swartpuntia* displaying no significant associations with either other soft-bodied taxa, nor biomineralisers while *Rangea* and *Pteridium* showed significant positive association with each other and negative associations with *Cloudina*. As such, there are no consistent patterns of biomineralisers nor soft-bodied taxa that explain the patterns found within our data.

As such, the signal in our data cannot be attributed solely to carbonate/siliclastic nor biomineralisers/soft-bodied taxa differences, and so is most likely due to habitat preferences as community composition was found to vary significantly with depth *Cloudina* is found exclusively in shallow limestone and shallow siliciclastic shoreface facies, whereas soft-bodied Nama taxa are found in both deeper shoreface and deep subtidal settings (Fig. 4).

**Figure 4.**
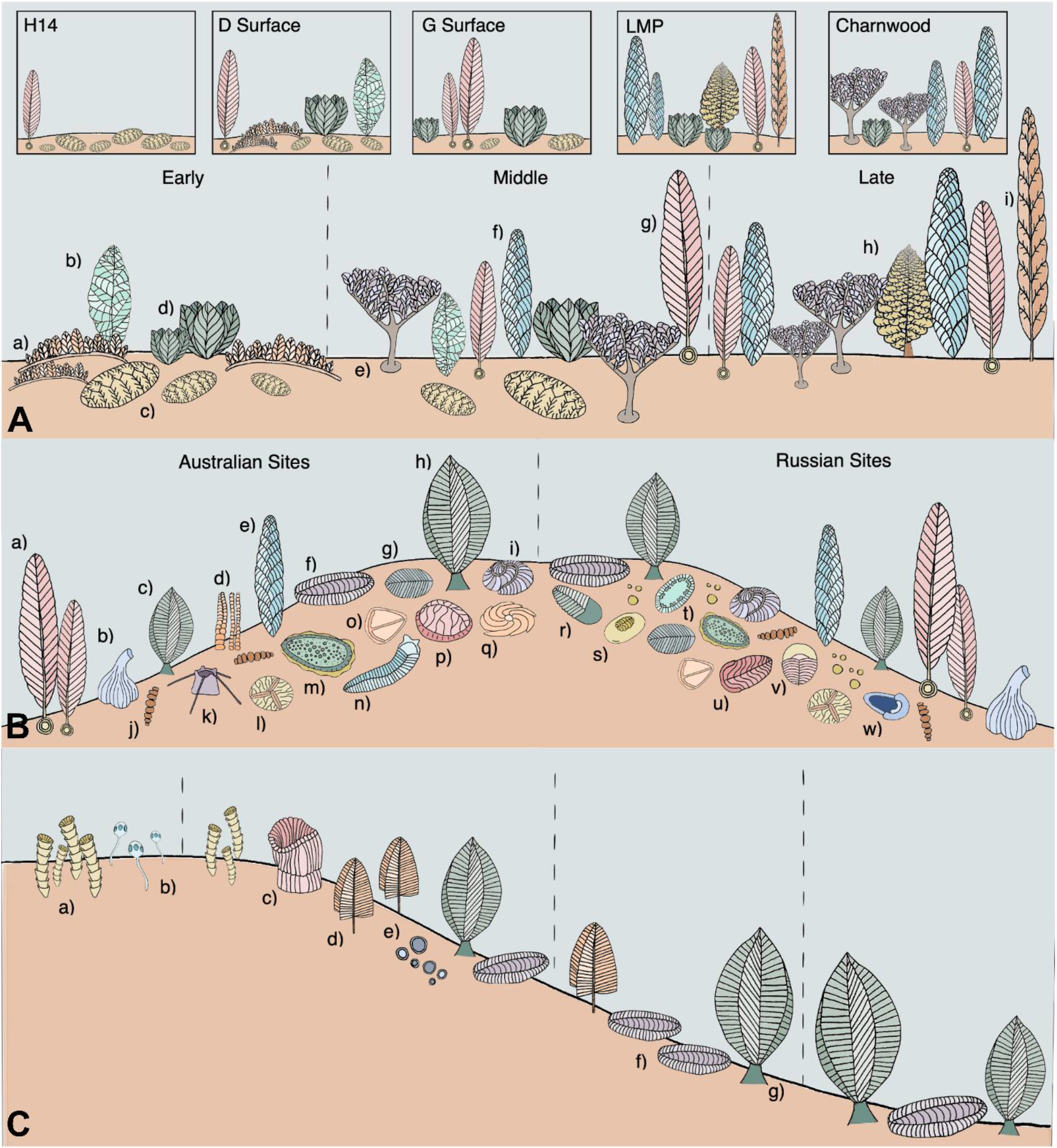
Reconstructions of the Avalon, White Sea and Nama metacommunities. **a**, A reconstruction of the Avalon assemblage showing the proposed stages of a community succession with the actual composition of several surfaces in boxes above. a) *Pectinifrons*, b) *Beothukis*, c) *Fractofusus*, d) *Bradgatia*, e) *Primocandelabrum*, f) *Charnia*, g) *Charniodiscus*, h) *Culmofrons*, i) *Trepassia*. **b**, A reconstruction of the White Sea assemblage showing some endemism of taxa to the Russian or Australian sites. a) *Charniodiscus*, b) *Inaria*, c) *Rangea*, d) *Funisia*, e) *Charnia*, f) *Pteridinium*, g) *Dickinsonia*, h) *Rangea*, i) *Tribrachidium*, j) *Palaeopaschinus*, k) *Coronacollina*, l) *Albumares*, m) *Kimberella*, n) *Spriggina*, o) *Parvancorina*, p) *Rugoconites*, q) *Eoandromeda*, r) *Cyanorus*, s) *Onega*, t) *Armillifera*, u) *Andiva*, v) *Yorgia*, w) *Temnoxa*. **c**, A reconstruction of the Nama assemblage showing the palaeoenvironmental separation of biomineralizing and soft-bodied taxa across a depth profile. a) *Cloudina*, b) *Namacalthus*, c) *Ernietta*, d) *Swartpuntia*, e) *Nimbia*, f) *Pteridinium*, g) *Rangea*. Taxa and environmental separation are not to scale.

## Discussion

Avalonian assemblage, with a ‘second-wave’ radiation encompassing the White Sea assemblage. A final radiation is observed in the Nama assemblage (5,29,30). The second-wave includes the development of the bilaterian body-plan, grazing, herbivory and wide-spread motility (29,30). These innovations are coupled to the development of dense communities with high community heterogeneity between environments (30,31) and increased taxa sensitivity to fine-scale environment (15,18). The second-wave is reflected in our results in an increase of non-random taxa associations from the Avalonian biota (9.8%) to the White Sea biota (16.1%). Further evidence for this increased ecological complexity is provided by the fact that the Avalonian assemblage had minimal environmental structuring, whilst we detected both Gleasonian and quasi-Clementsian metacommunity structure depending on the palaeoenvironments for the White Sea assemblage. While there is not a significant correlation of broad-scale palaeoenvironment with metacommunity structure, these structures reflect a significant influence of a fine-scale environmental gradient. Gleasonian structuring reflects an individualistic response to the inferred environmental gradient, suggesting a lack of within-community interactions for this outer shelf metacommunity. In contrast, quasi-Clementsian structure corresponds to a community-wide response to the environmental gradients, so reflects within-community specialisations in the White Sea assemblage. These specialisations are reflected in behavioral flexibility, with organisms acting differently in different environments across a wide range of depths.

The White Sea and Nama assemblages have the same type of metacommunity structure as shown through EMS, with both assemblages showing a quasi-Clementsian structure (Fig. 3, Table 1). There is a small increase in non-random associations in the Nama biota (16.7% from 16.1% in the White Sea), which, like the EMS analyses shows at least a maintenance if not slight increase in metacommunity structure. There is a significant increase in ecosystem structuring between the White Sea and Nama assemblages when site rank within each of the assemblages are compared to depth (Table 2). Neither the Avalon nor White Sea showing significant correlation in sharp contrast to the Nama, where site composition is significantly correlated with depth (Table 2). Taken together, these three analyses show that compared to the White Sea, the Nama assemblage has an increased taxa segregation coupled to a strong palaeoenvironmental correlation and therefore narrowed environmental tolerances, showing a further decrease of niche breadth. Thus, we have shown that the increase in complexity of the autecological strategies utilised throughout the Ediacaran is mirrored in the complexity of the synecological interactions.

The reduction in taxonomic diversity and community composition complexity has been suggested to correspond to a post-White Sea extinction caused by either an environmental driven catastrophic environmental extinction or biotic replacement driven extinction (12,23,22,24,21,21). Recent work has shown that a biotic replacement driven extinction, whereby mobile metazoans out-competed soft-bodied Ediacaran organisms through bioturbation and ecosystem engineering, is unlikely due in part to prolonged co-occurrence of trace fossils with soft-bodied biota (5,32). Other currently unknown and/or unpreserved intrinsic causes behind a biotic replacement model cannot not be excluded at the moment. A White Sea – Nama catastrophic environmental extinction implies that surviving taxa within the Nama assemblage were generalists (11,12,33), contrary to our results. We have shown an increased influence of paleoenvironment and niche specialisation with a change from nested to Clementsian structuring indicating more Turnover along the gradient (more niche differentiation). If the Nama assemblage metacommunity structure was due to underlying extinction/colonization dynamics, we would expected to see an increase of nestedness (34), as indicated by negative Turnover, contrary to our positive Nama Turnover (Table 1). This increase in Turnover suggests not only higher ecosystem complexity but also increased taxa specialisation and narrower niche breadths is coupled with an increase in within-community structuring between the White Sea and Nama assemblages, with an increase in non-random taxa associations (16.1% to 16.7%). A decrease in taxonomic and morphological diversity in the Nama (23) may reflect that, within this assemblage, Ediacaran organisms show significant palaeoenvironmental preferences, and thus reduced environmental tolerances, resulting in multiple different types of mutually exclusive communities, each of which exhibits a simple structure within its narrow niche (23). An increase in within-community structure in the form of ecosystem engineering (35) and reef complexity (20) provides supporting evidence that despite a decrease in taxonomic diversity, the Nama assemblage represents an ecological development from the White Sea assemblage, not a recovery from an catastrophic extinction event. Our results are further supported by birth-death models of stem and crown group diversification, which predict Ediacaran-like diversification patterns for bilaterians and produce patterns which can be easily mistaken for mass extinctions (36).

Our results show that these Ediacaran organisms underwent the niche contraction and specialisation that is traditionally associated with Cambrian diversification (9,37). Therefore, we find that the eco-evolutionary dynamics of metazoan diversification known from the Cambrian started earlier in the Ediacaran with the Avalon assemblage and increased in complexity towards the Phanerozoic as new anatomical innovations appeared, culminating in the “Cambrian Explosion”.

## Materials and Methods

### Materials

Ediacaran fossils are commonly found preserved in-situ, so their bedding-planes preserve near-compete censuses of the communities (29,38,38). This exceptional preservation means that ecological analyses normally reserved for modern communities can be applied e.g. (15,39). The data used in this study is a binary presence/absence matrix for 86 Ediacaran localities and 124 taxa. The data is taken from (12) with more conservative classifications of assemblages for several sites (cf. (11)): SB-Nor2 and SB-So1, both Sewki Brook sites, are classed as indeterminate in our analysis and (11) but as Avalon and Nama respectively in (12). Two Chinese sites classed as Nama by Muscente et al. are classed as Indeterminate here (Gaojiashan and Lijiagou) (11,24). The data contains information on the palaeoenvironment, depth index (from 1-11), lithology and assemblage of each locality as well as a time index (from 1-3). The full classification of sites in the dataset can be seen in Data S1 and the palaeoenvironmental and assemblage classifications in Fig. 1. This data is appropriate for applying modern statistical ecological methods because the organisms were mostly sessile and benthic and preserved in such a way that they are interpreted as in-situ life assemblages with minimal transportation after death or time-averaging (16,25,38,39).

The Avalon, White Sea and Nama assemblages are represented by 30, 29 and 12 sites respectively, with 11 undetermined sites and 4 which belong to the Doushantuo assemblage. The Doushantuo assemblage was excluded from the assemblage-level analyses because there are only four sites, which is insufficient to run these analyses.

#### Elements of Metacommunity Structure (EMS) analysis

Three metacommunity metrics are calculated to determine the metacommunity structure (Fig. 1). First, Coherence is a measure of the extent to which all the taxa respond to the same environmental gradient, where this gradient may result from the interplay of several biotic and abiotic factors that differ between sites (1). Coherence is positive when the taxa in the site-by-taxa matrix all respond to the same environmental gradient. Most extant well-sampled metacommunities display significant positive Coherence due to similarities in evolutionary history, ecological preferences or life-history trade-offs within communities (1). A significantly negatively Coherent site-by-taxa matrix, reflects a high number of mutually exclusive taxa pairs creating checkerboard patterns (10,12,13). These checkerboard patterns are random with respect to each other and there is no discernable gradient to which all the taxa respond. Negative co-occurrences and significant segregation/checkerboard patterns can be formed from strong competition, grazing/herbivory or strongly non-overlapping niches, all of which form similar metacommunity patterns due the presence of mutually exclusive pairs of taxa (1,2,4). A non-significant Coherence reflects no significant metacommunity structuring. For metacommunities which have positive Coherence, the Turnover metric tests the amount of taxa replacement between sites (40). If taxa ranges are nested within each other, there is less Turnover than expected by chance along the gradient (significant negative). If there are more differences in site taxa composition along the gradient than expected by chance, Turnover is significantly positive and the structure is non-nested. Non-significant Turnover indicates a weaker structuring mechanism, a quasi-structure (3). The final metric, Boundary Clumping measures the extent to which taxa range limits cluster at the same sites across the environmental gradient (1). The range limits can be clumped (significant positive), hyperdispersed (significant negative) or random (non-significant).

Positive Coherence and negative Turnover results in nested metacommunities with taxa-poor sites being predictable sub-sets of taxa-rich sites. Depending on Boundary Clumping, taxa loss can be hyper-dispersed, random or clumped (16). Positive Coherence and Turnover with negative Boundary Clumping describes evenly spaced metacommunity. Where Coherence, Turnover and Boundary Clumping are all positive, the metacommunity is classed as Clementsian (Fig. 1), where groups of taxa with similar range boundaries co-occur and respond in a similar way to environment gradients (1,41). Taxa within Clementsian metacommunities respond synchronously to environmental gradients, suggesting physiological or evolutionary trade-offs associated with environmental thresholds (42). When Coherence and Turnover are positive but there is no significant Boundary Clumping, the metacommunity is described as Gleasonian where each taxa reacts individualistically to environmental gradients (3).

The R package Metacom was used to quantify the three metrics related to metacommunity structure: Coherence, Turnover and Boundary Clumping (1). A three-tiered analysis based on these metrics enabled the placement of each studied metacommunity into one of 12 idealized metacommunity structures following the Elements of Metacommunity Structure approach (EMS) (2,3). Significant Coherence is a pre-requisite for further analysis of metacommunity structure and so this was the first metric calculated. Calculation of Turnover was then used to distinguish whether the metacommunity formed a nested structure. Calculation of Boundary Clumping allowed the determination of whether the taxa range boundaries were clumped, dispersed or not significantly correlated with each other along the environmental gradient. Figure 1 gives examples of taxa abundance distributions that give rise to the idealized metacommunity structures. Sites scores were given according to the ranking of the sites in the first-degree ordination of their taxa composition. These were used to investigate the importance of depth (as indicated by palaeoenvironment) and assemblage variables in structuring taxa distributions via a non-parametric Spearman’s correlation or Kruskal-Wallis test. The metacommunity analyses were performed on the entire dataset, for each assemblage individually, and for palaeoenvironmental and geographic subsets within each assemblage where there were enough localities for valid analyses.

#### Co-occurrence analysis

The R package cooccur was used to calculate the observed and expected frequency of co-occurrence between pairs of taxa to determine significant positive or negative associations (42). Taxa which only occurred in one site were removed from the analysis because such singletons have been shown to disproportionally influence co-occurrence analyses (43,44). Co-occurrence analysis was done for the entire dataset, for each assemblage individually, and for palaeoenvironmental and geographic subsets within each assemblage where there were enough sites (42).

## Funding

Natural Environment Research Council Independent Research Fellowship NE/S014756/1 to EGM. The funders had no role in study design, data collection and analysis, decision to publish, or preparation of the manuscript.

## Author contributions

EGM conceived the initial study with input from AM and all authors developed the work. RE performed the analyses. All authors discussed the results and wrote the manuscript.

## Competing interests

Authors declare no competing interests.

## Data and materials availability

All data and code is available in the supplementary materials. The data used in this paper has been modified from previously published data and will be made publicly available on figshare doi: 10.6084/m9.figshare.13664105 (private link before doi is published: https://figshare.com/s/52bc12af68d151663ee2)

## Supplementary Information

Supplementary discussion

Supplementary Figures S1-4.

Supplementary Tables S1-9.

## Notes

### Competing Interest Statement

The authors have declared no competing interest.

## References

1. Leibold M A, Mikkelson G M. Coherence, species turnover, and boundary clumping: elements of meta-community structure. Oikos [Internet]. 2002 Jul 1 [cited 2020 Mar 1];97(2). Available from: https://onlinelibrary.wiley.com/doi/epdf/10.1034/j.1600-0706.2002.970210.x

2. Leibold MA, Holyoak M, Mouquet N, Amarasekare P, Chase JM, Hoopes MF, et al. The metacommunity concept: a framework for multi-scale community ecology. Ecology Letters. 2004;7(7):601–13.

3. Presley S, Higgins CL, Willig MR. A comprehensive framework for the evaluation of metacommunity structure. Oikos. 2010;119(6):908–17.

4. Dallas T. metacom: an R package for the analysis of metacommunity structure. Ecography. 2014;37(4):402–5.

5. Wood R, Liu AG, Bowyer F, Wilby PR, Dunn FS, Kenchington CG, et al. Integrated records of environmental change and evolution challenge the Cambrian Explosion. Nat Ecol Evol. 2019;3(4):528–38.

6. Bobrovskiy I, Hope JM, Krasnova A, Ivantsov A, Brocks JJ. Molecular fossils from organically preserved Ediacara biota reveal cyanobacterial origin for Beltanelliformis. Nature Ecology & Evolution. 2018 Mar;2(3):437–40.

7. Xiao S, Laflamme M. On the eve of animal radiation: phylogeny, ecology and evolution of the Ediacara biota. Trends in Ecology and Evolution. 2009;24(1):31–40.

8. Shen B, Dong L, Xiao S, Kowalewski M. The Avalon Explosion: Evolution of Ediacara Morphospace. Science. 2008 Jan 4;319(5859):81–4.

9. Bush AM, Bambach RK, Erwin DH. Ecospace Utilization During the Ediacaran Radiation and the Cambrian Eco-explosion. Quantifying the Evolution of Early Life. 2011;111–33.

10. Waggoner B. The Ediacaran Biotas in Space and Time. Integr Comp Biol. 2003 Feb 1;43(1):104– 13.

11. Boag TH, Darroch SAF, Laflamme M. Ediacaran distributions in space and time: testing assemblage concepts of earliest macroscopic body fossils. Paleobiology. 2016;42(4):574–94.

12. Muscente AD, Bykova N, Boag TH, Buatois LA, Mángano MG, Eleish A, et al. Ediacaran biozones identified with network analysis provide evidence for pulsed extinctions of early complex life. Nat Commun [Internet]. 2019 Feb 22 [cited 2019 Oct 14];10. Available from: https://www.ncbi.nlm.nih.gov/pmc/articles/PMC6384941/

13. Grazhdankin D. Patterns of distribution in the Ediacaran biotas: facies versus biogeography and evolution. Paleobiology. 2004 Mar 1;30(2):203–21.

14. Mitchell EG, Butterfield NJ. Spatial analyses of Ediacaran communities at Mistaken Point. Paleobiology. 2018 Feb 1;44(1):40–57.

15. Mitchell EG, Harris S, Kenchington CG, Vixseboxse P, Roberts L, Clark C, et al. The importance of neutral over niche processes in structuring Ediacaran early animal communities. Ecology Letters. 2019;22(12):2028–38.

16. Clapham ME, Narbonne GM, Gehling JG. Paleoecology of the Oldest Known Animal Communities: Ediacaran Assemblages at Mistaken Point, Newfoundland. Paleobiology. 2003;29(4):527–44.

17. Mitchell EG, Kenchington CG. The utility of height for the Ediacaran organisms of Mistaken Point. Nat Ecol Evol. 2018 Jun 25;2:1218–22.

18. Mitchell EG, Bobkov N, Bykova N, Dhungana A, Kolesnikov A, Hogarth IRP, et al. The influence of environmental setting on the community ecology of Ediacaran organisms. Interface Focus [Internet]. 2020.[cited 2020 May 17]; Available from: https://www.biorxiv.org/content/10.1101/861906v1

19. Narbonne GM, Gehling JG. Life after snowball: The oldest complex Ediacaran fossils. Geology. 2003 Jan 1;31(1):27–30.

20. Penny AM, Wood R, Curtis A, Bowyer F, Tostevin R, Hoffman K-H. Ediacaran metazoan reefs from the Nama Group, Namibia. Science. 2014;344(6191):1504–6.

21. Darroch SAF, Sperling EA, Boag TH, Racicot RA, Mason SJ, Morgan AS, et al. Biotic replacement and mass extinction of the Ediacara biota. Proceedings of the Royal Society B: Biological Sciences. 2015 Sep 7;282(1814):20151003.

22. Darroch SAF, Smith EF, Laflamme M, Erwin DH. Ediacaran Extinction and Cambrian Explosion. Trends in Ecology & Evolution. 2018 Sep 1;33(9):653–63.

23. Darroch SAF, Laflamme M, Wagner PJ. High ecological complexity in benthic Ediacaran communities. Nature Ecology & Evolution. 2018 Oct;2(10):1541–7.

24. Muscente AD, Boag TH, Bykova N, Schiffbauer JD. Environmental disturbance, resource availability, and biologic turnover at the dawn of animal life. Earth-Science Reviews. 2018 Feb 1;177:248–64.

25. Droser ML, Gehling JG, Jensen SR. Assemblage palaeoecology of the Ediacara biota: The unabridged edition? Palaeogeography, Palaeoclimatology, Palaeoecology. 2006 Mar 22;232(2):131–47.

26. García-Girón J, Heino J, Alahuhta J, Chiarenza AA, Brusatte SL. Palaeontology meets metacommunity ecology: the Maastrichtian dinosaur fossil record of North America as a case study. Palaeontology. 2021;1–23.

27. Bar-Massada A. Complex relationships between species niches and environmental heterogeneity affect species co-occurrence patterns in modelled and real communities. Proceedings of the Royal Society B: Biological Sciences. 2015 Aug 22;282(1813):20150927.

28. Grazhdankin D, Gerdes G. Ediacaran microbial colonies. Lethaia. 2007;40(3):201–10.

29. Droser ML, Gehling JG. The advent of animals: The view from the Ediacaran. PNAS. 2015 Apr 21;112(16):4865–70.

30. Droser ML, Gehling JG, Tarhan LG, Evans SD, Hall CMS, Hughes IV, et al. Piecing together the puzzle of the Ediacara Biota: Excavation and reconstruction at the Ediacara National Heritage site Nilpena (South Australia). Palaeogeography, Palaeoclimatology, Palaeoecology. 2019 Jan 1;513:132–45.

31. Finnegan S, Gehling JG, Droser ML. Unusually variable paleocommunity composition in the oldest metazoan fossil assemblages. Paleobiology. 2019 May;45(2):235–45.

32. Darroch SAF, Cribb AT, Buatois LA, Germs GJB, Kenchington CG, Smith EF, et al. The trace fossil record of the Nama Group, Namibia: Exploring the terminal Ediacaran roots of the Cambrian explosion. Earth-Science Reviews. 2021 Jan 1;212:103435.

33. Schiffbauer JD, Huntley JW, O’Neil GR, Darroch SAF, Laflamme M, Cai Y. The Latest Ediacaran Wormworld Fauna: Setting the Ecological Stage for the Cambrian Explosion. GSAT. 2016 Nov 1;4–11.

34. Wright DH, Patterson BD, Mikkelson GM, Cutler A, Atmar W. A Comparative Analysis of Nested Subset Patterns of Species Composition. Oecologia. 1998;113(1):1–20.

35. Cribb AT, Kenchington CG, Koester B, Gibson BM, Boag TH, Racicot RA, et al. Increase in metazoan ecosystem engineering prior to the Ediacaran–Cambrian boundary in the Nama Group, Namibia. Royal Society Open Science. 2019;6(9):190548.

36. Budd GE, Mann RP. The dynamics of stem and crown groups. Science Advances. 2020 Feb 1;6(8):eaaz1626.

37. Na L, Kiessling W. Diversity partitioning during the Cambrian radiation. PNAS. 2015 Apr 14;112(15):4702–6.

38. Wood DA, Dalrymple RW, Narbonne GM, Gehling JG, Clapham ME. Paleoenvironmental analysis of the late Neoproterozoic Mistaken Point and Trepassey formations, southeastern Newfoundland. Canadian Journal of Earth Sciences. 2003 Oct 1;40(10):1375–91.

39. Mitchell EG, Kenchington CG, Liu AG, Matthews JJ, Butterfield NJ. Reconstructing the reproductive mode of an Ediacaran macro-organism. Nature. 2015 Aug;524(7565):343–6.

40. Patterson BD, Atmar W. Nested subsets and the structure of insular mammalian faunas and archipelagos. Biol J Linn Soc. 1986 May 1;28(1–2):65–82.

41. Dahlgren. Incorporating environmental change over succession in an integral projection model of population dynamics of a forest herb. Oikos. 2011;20:1183–90.

42. Griffith DM, Veech JA, Marsh CJ. cooccur: Probabilistic Species Co-Occurrence Analysis in R. Journal of Statistical Software. 2016 Feb 9;69(1):1–17.

43. Collins MD, Simberloff D, Connor EF. Binary matrices and checkerboard distributions of birds in the Bismarck Archipelago. Journal of Biogeography. 2011;38(12):2373–83.

44. Pitta E, Giokas S, Sfenthourakis S. Significant Pairwise Co-occurrence Patterns Are Not the Rule in the Majority of Biotic Communities. Diversity. 2012 Jun;4(2):179–93.

